# Defined diets for freshwater planarians

**DOI:** 10.1101/2021.04.05.438509

**Authors:** Chris Abel, Kaleigh Powers, Gargi Gurung, Jason Pellettieri

## Abstract

Planarian flatworms are popular invertebrate models for basic research on stem cell biology and regeneration. These animals are commonly maintained on a diet of homogenized calf liver or boiled egg yolk in the laboratory, introducing a source of uncontrolled experimental variability. Here, we report the development of defined diets, prepared entirely from standardized, commercially sourced ingredients, for the freshwater species *Schmidtea mediterranea, Dugesia japonica*, and *Girardia dorotocephala*. These food sources provide an opportunity to test the effects of specific nutritional variables on biological phenomena of interest. Defined diet consumption was not sufficient for growth and only partially induced the increase in stem cell division that normally accompanies feeding, suggesting these responses are not solely determined by caloric intake. While our defined diet formulations do not support long-term planarian maintenance, they do enable delivery of double-stranded RNA for gene knockdown in a manner that provides unique advantages in some experimental contexts. We also present a new approach for preserving tissue integrity during hydrogen peroxide bleaching of liver-fed animals. These tools will empower research on the connections between diet, metabolism, and stem cell biology in the experimentally tractable planarian system.

## INTRODUCTION

Freshwater planarians have long been recognized for their remarkable ability to regenerate any lost body part, and the recent application of experimental tools like RNA interference (RNAi) and RNA sequencing has led to a better mechanistic understanding of this process (Newmark & Sánchez Alvarado, 2002; Ivankovic et al., 2019). Most notably, we now know far more about the adult stem cells, or ‘neoblasts,’ that drive formation of new tissue at sites of amputation (Reddien, 2018; Dattani et al., 2019). Under homeostatic conditions, these cells remain mitotically active to enable a high rate of physiological cell turnover. Following injury, they increase their rate of division and migrate to the wound site, where they give rise to a mass of undifferentiated tissue called the blastema that subsequently differentiates to restore a complete and functional anatomy.

Neoblasts are also responsive to nutritional cues, increasing their rate of division within hours of feeding and remaining in a state of heightened proliferation for up to a few days (Baguñà, 1974; Baguñà, 1976; Baguñà & Romero, 1981; Salò and Baguñà, 1984; Newmark & Sánchez Alvarado, 2000; Kang & Sánchez Alvarado, 2009; González-Estévez et al., 2012; Pascual-Carreras et al., 2020). In asexual strains of freshwater species like *S. mediterranea, D. japonica*, and *G. dorotocephala*, accompanying organismal growth eventually leads to reproduction via binary fission (Vila-Farré & Rink, 2018). During prolonged starvation, planarians undergo a more than 40-fold reduction in overall size, a process historically termed ‘degrowth’ (Baguñà et al., 1990; Newmark & Sánchez Alvarado, 2002). This response is primarily attributed to a change in cell number, rather than a change in cell size – neoblasts do not alter their rate of mitosis, but their differentiating division progeny decrease in number while the rate of cell death rises (Baguñà, 1976; Pellettieri et al., 2010; González-Estévez et al., 2012; Thommen et al., 2019). Animals retain anatomical scale and proportion as they shrink, both at the organismal level and with respect to specific differentiated tissues and cell types (Oviedo et al., 2003; Takeda et al., 2009; Pellettieri et al., 2010; Forsthoefel et al., 2011). These observations point to the existence of mechanisms coordinating stem cell division, cell death, and organismal growth. Although recent studies have revealed some of the genetic pathways that control these processes (Oviedo et al., 2008a; Bender et al., 2012; Miller & Newmark, 2012; Peiris et al., 2012; Tu et al., 2012; Almuedo-Castillo et al., 2014; Lin & Pearson, 2014; Lin & Pearson, 2017; de Sousa et al., 2018; Arnold et al., 2019; Pascual-Carreras et al., 2020; Schad & Petersen, 2020; Ziman et al., 2020), it is not yet clear how cell number is “counted” or attuned to metabolic cues to effect rapid transitions between growth and degrowth.

Planarians are carnivores. In the wild, they are reported to consume living or recently dead arthropods (e.g., insect larvae and crustaceans), annelids (e.g., oligochaetes), and molluscs (e.g., gastropods) (Jennings, 1957; Reynoldson & Young, 1963; Pickavance, 1971; Boddington & Mettrick, 1974; Reynoldson & Sefton, 1976; Armitage & Young, 1990; Gee & Young, 1993; Calow et al., 2009; Vila-Farré & Rink, 2018; Cuevas-Caballé et al., 2019). Some species may use their mucus secretions to entrap prey (Jennings, 1957). Once a food item is encountered, the cylindrical pharynx is extended through a ventral opening in the body wall. The peristaltic activity of the pharyngeal muscles then draws tissues and body fluids from the animal on which the planarian is feeding into the highly branched gut, where absorptive phagocytes internalize and break down the material to fuel anabolic processes (Ishii & Sakurai, 1991; Adler et al., 2014; Forsthoeffel et al., 2020). Undigested food is eliminated through the pharynx, while a protonephridial system allows for excretion of fluid waste (Hertel, 1993; Rink et al., 2011; Scimone et al., 2011).

Freshwater planarians used in stem cell and regeneration research are typically maintained on a diet of homogenized calf liver or boiled egg yolk in the laboratory (Oviedo et al., 2008b; Accorsi et al., 2017; Merryman et al., 2018). Although these diets are inexpensive, relatively easy to prepare, and promote rapid growth, their lack of standardization introduces an uncontrolled source of variability, both within and between research groups. This is a particular concern in studies addressing links between nutrition and physiology. Furthermore, such non-defined (oligidic) diets afford only limited potential for experimental manipulation of nutritional variables. To address these concerns, we have developed meridic diets, prepared almost entirely from chemically defined ingredients, and used them to begin analyzing the nutritional basis for the mitogenic effect of feeding. We show that defined diet consumption is sufficient for partial induction of neoblast proliferation, but not to support overall organismal growth. Thus, like defined diets previously developed for *Drosophila melanogaster* and *Caenorhabditis elegans*, the formulations we describe here for planarians are not optimal for long-term propagation of laboratory populations (Lüersen et al., 2019; Zečić et al., 2019).

Nevertheless, we expect them to prove similarly useful in empowering research on the relationships between nutrition, metabolism, and the biological phenomena for which these experimentally tractable systems are so well known.

## RESULTS AND DISCUSSION

### Development and characterization of defined diets

In an effort to develop a simple, standardized food source for freshwater planarians, we prepared defined diets from commercially sourced ingredients and tested their ability to stimulate feeding behavior. Our formulations were based upon a nutritional analysis of beef liver (Haytowitz et al., 2019), with adjustments to the concentrations of some ingredients to maintain solubility or promote feeding. Through this empirical approach, we arrived at two diets, hereafter referred to as complex and minimal. The former consists of the 20 standard amino acids, dextrose, glycogen, lecithin, and xanthan gum as a thickener; the latter consists only of lecithin and xanthan gum (Table 1; Materials and Methods).

**Table 1.** Defined diet formulations. See Supplementary Materials and Methods for detailed protocols on the preparation of each diet.

| **Compound** | **Vendor** | **Catalog No.** | **g per 10 ml** |
| --- | --- | --- | --- |
| L-Valine | Acros Organics | 140810250 | 0.015 |
| L-Alanine | Acros Organics | 102831000 | 0.016 |
| L-Leucine | Sigma-Aldrich | L8912 | 0.022 |
| L-Methionine | Sigma-Aldrich | M9625 | 0.006 |
| L-Isoleucine | Acros Organics | 166170250 | 0.011 |
| L-Cysteine | Sigma-Aldrich | C7352 | 0.004 |
| L-Tyrosine | Sigma-Aldrich | T3754 | 0.001 |
| L-Phenylalanine | Acros Organics | 130311000 | 0.014 |
| L-Threonine | Acros Organics | 138930250 | 0.011 |
| L-Aspartic acid | Fisher Scientific | BP374 | 0.026 |
| L-Glutamic acid, monosodium salt hydrate | Sigma-Aldrich | G5889 | 0.045 |
| Glycine | Sigma-Aldrich | G7126 | 0.022 |
| L-Proline | Sigma-Aldrich | P0380 | 0.016 |
| L-Serine | Acros Organics | 132661000 | 0.013 |
| L-Glutamine | Sigma-Aldrich | G3126 | 0.004 |
| L-Histidine | Acros Organics | 166150250 | 0.008 |
| L-Lysine monohydrochloride | Alfa Aesar | J62099 | 0.026 |
| L-Arginine monohydrochloride | Sigma-Aldrich | A5131 | 0.022 |
| L-Tryptophan | Acros Organics | 140590250 | 0.003 |
| L-Asparagine | Acros Organics | 371601000 | 0.003 |
| Dextrose | Fisher Scientific | D16 | 0.022 |
| Glycogen | BioBasic | GB0301 | 0.300 |
| Egg yolk lecithin | Sigma-Aldrich | P5394 | 0.700 |
| Xanthan gum | Sigma-Aldrich | G1253 | 0.025 |

**MINIMAL DIET**
| **Compound** | **Vendor** | **Catalog No.** | **g per 10 ml** |
| --- | --- | --- | --- |
| Egg yolk lecithin | Sigma-Aldrich | P5394 | 0.700 |
| Xanthan gum | Sigma-Aldrich | G1253 | 0.025 |

Starved *S. mediterranea, D. japonica*, and *G. dorotocephala* all consumed both the complex and minimal diets under ad libitum feeding conditions (Materials and Methods), though the percentage of animals that ate and the amount of food consumed was not as high as for liver (Figure 1; Figure S1). Because it was particularly difficult to induce long-term feeding with the minimal diet (Figure 1B), our use of this formulation in follow-up experiments was restricted to those cases in which the effects of a single feeding were analyzed. Together, the complex and minimal diets allow for complementary approaches to studying the impacts of nutritional cues on planarian biology – ingredients can be subtracted from the complex diet in order to determine whether they are necessary for phenomena of interest and/or added to the minimal diet in order to determine whether they are sufficient. For example, supplementing the minimal diet (or xanthan gum alone) may represent a useful strategy for identifying additional chemoattractants that, like lecithin, stimulate feeding behavior.

**Figure 1.**
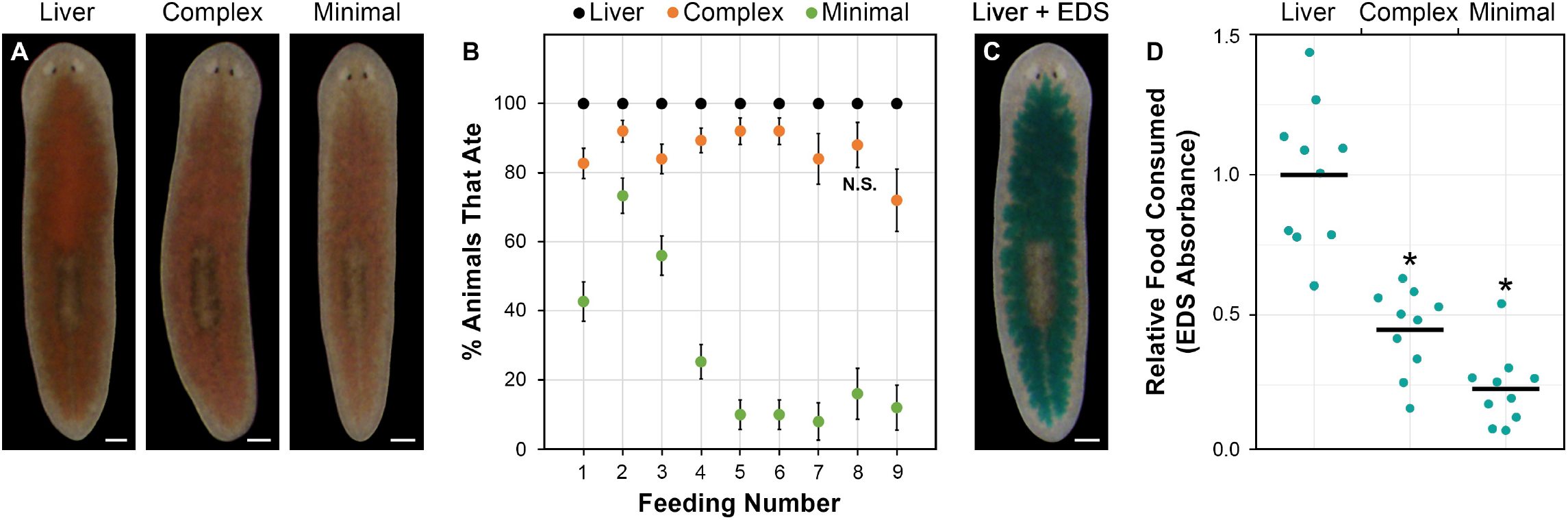
Defined diet feeding in *S. mediterranea*. **(A)** Representative images of live animals photographed after a single ad libitum feeding with homogenized beef liver or defined diets. Red food coloring was added to visualize ingested food. **(B)** Percentage of animals that consumed each diet. Results are from three biological replicates, each of which included 25 animals per condition. Feedings were administered every other day until the liver-fed animals began to fission (replicates were terminated after feedings 4, 6, and 9). Error bars show standard error. Differences between liver and each defined diet were significant at all time points (Pearson’s chi-square test; p < 0.05), except for the complex diet in feeding number 8. **(C)** Representative image of an animal fed liver containing 0.2% erioglaucine disodium salt (EDS). **(D)** The relative amount of each diet consumed in a single ad libitum feeding was determined by measuring EDS absorbance in lysates of fed animals (Materials and Methods). Each data point denotes the absorbance of a single lysate prepared from five animals, with all values expressed relative to the mean for liver. Horizontal lines denote means. Asterisks denote p values < 5.0E-05 for T-tests comparing liver and each defined diet. Scale bars: (A,C) = 200 µm.

### Growth and stem cell proliferation are not solely determined by caloric intake

To assess whether the complex diet could substitute for liver in the long-term laboratory propagation of *S. mediterranea*, we compared the ability of these two food sources to induce overall growth in size-matched animals. While feeding every other day with liver led to a rapid increase in animal area, followed by fission, animals fed on the same schedule with the complex diet displayed a gradual reduction in size (Figure 2A). This decline was not as pronounced as that observed in starved controls, however, indicating the complex diet has at least some nutritional value. 20% of animals maintained on the complex diet began to develop lesions and lyse beyond 25 feedings (50 days). This was not observed in the starved controls, potentially indicating that specific ingredients or altered nutrient ratios in the complex diet suppress catabolic processes necessary for survival or become toxic in the context of degrowth (Wulzen & Bahrs, 1931). We also evaluated whether our defined diet formulations could trigger the increase in stem cell division that normally occurs after feeding, using phosphorylated-histone H3 (H3P) immunostaining (Newmark & Sánchez Alvarado, 2000) (Materials and Methods). Liver consumption resulted in a slightly more than two-fold increase in H3P+ cells 24 hours after a single feeding (Figure 2B,C). Consumption of the minimal or complex diets also induced stem cell division, but to a significantly lower extent.

**Figure 2.**
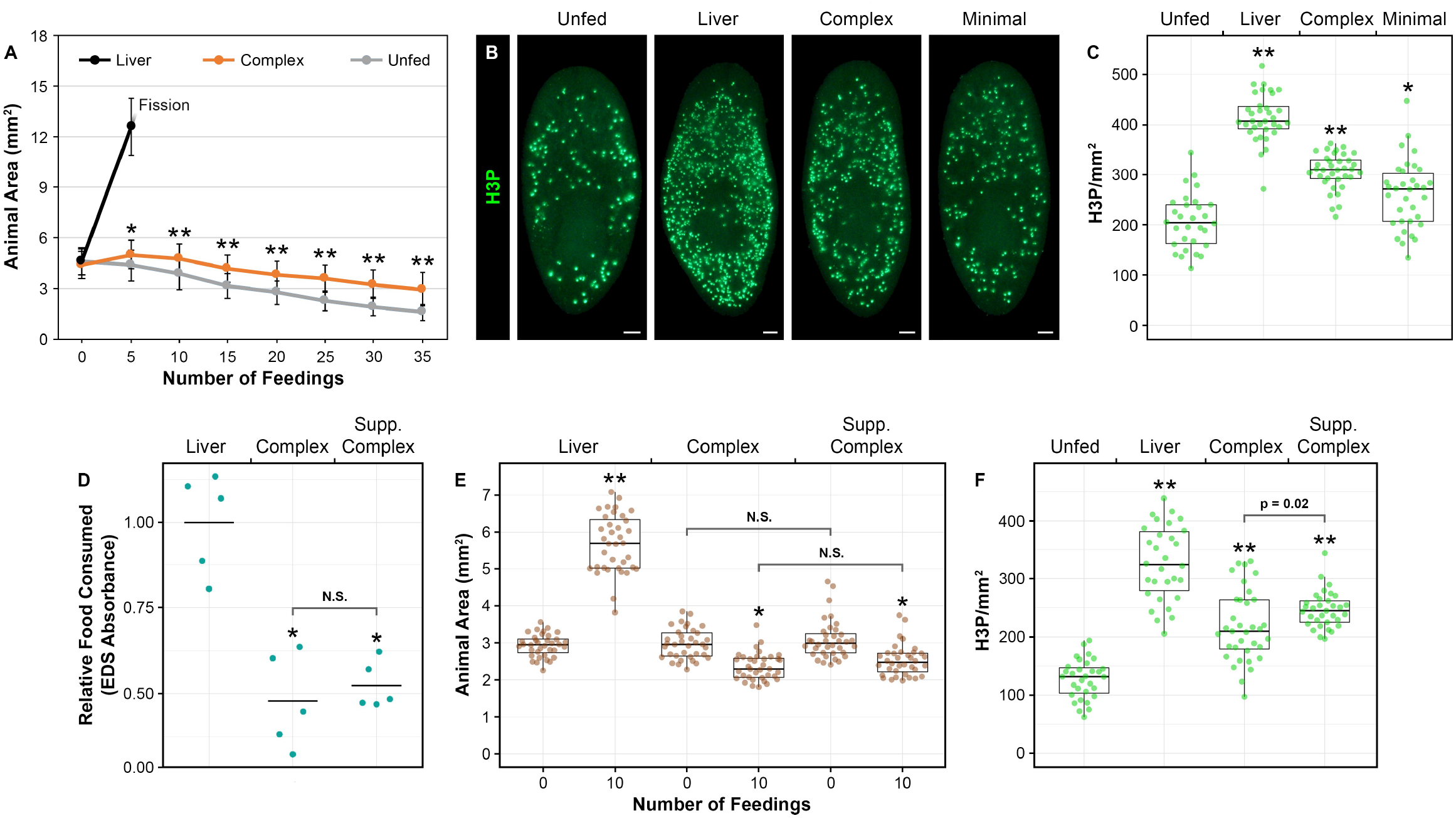
Impacts of defined diets on growth and stem cell division. **(A)** Ad libitum feeding with the complex diet every other day significantly slowed the rate of degrowth. Results are from three biological replicates, each of which included 15 animals per condition. Error bars show standard deviation. Asterisks and double asterisks denote p values < 0.002 and < 5.0E-05, respectively, for T-tests comparing unfed controls and animals fed with the complex diet. Liver-fed controls started to fission beyond 5 feedings and were excluded from subsequent analysis. **(B)** Representative images of H3P immunostaining in unfed controls and animals fixed 24 hours after a single feeding with the indicated diets. Scale bars = 100 µm. **(C)** Quantitative analysis of H3P results. Data are from three biological replicates, with a minimum of 30 combined animals per condition. Boxes, whiskers, and horizontal lines denote the interquartile range (IQR), values within 1.5X the IQR, and medians, respectively. Asterisks denote p values for T-tests comparing unfed controls and animals fed with the indicated diets (single asterisk: p < 1E-3; double asterisks: p < 1E-10). **(D-F)** Effects of complex diet supplementation (0.5 g casein and 2.0 g dextrose per 10 ml) on rates of food consumption (D), changes in animal size (E), and stem cell division at 24 hours post-feeding (F). Relative food consumption was determined from EDS absorbance measurements (each data point indicates the absorbance of a single lysate prepared from five animals, with values expressed relative to the mean for liver). Results for animal area and H3P labeling are from three biological replicates, with a minimum of 30 combined animals per condition. Horizontal lines denote means in (D) and medians in (E,F). Boxes and whiskers denote the IQR and range of values within 1.5X the IQR, respectively. Asterisks denote T-test p values for comparisons with liver-fed controls (D), 0-feeding controls (E), or unfed controls (F), except for the indicated, direct comparisons between the standard and supplemented complex diet formulations (single asterisk: p < 1E-3; double asterisks: p < 5E-10; N.S.: p > 0.05).

The caloric content of beef liver is approximately 1.5 kcal/ml (1.4 kcal/g) (Haytowitz et al., 2019). By comparison, we estimated the minimal and complex diets contain only 0.7 and 0.9 kcal/ml, respectively, with their overall nutritional values being further diminished by their reduced rates of consumption (Figure 1D). Thus, we reasoned differences in growth/degrowth and feeding-induced neoblast division in the above experiments might simply be due to differences in total caloric intake. To address this possibility, we next tested the effect of doubling the caloric value of the complex diet via addition of 0.45 kcal/ml each casein and dextrose. This modification did not significantly impact the amount of food consumed (Figure 2D), indicating it also increased total caloric intake by approximately two-fold. Yet it still failed to induce growth, or even to slow the rate of degrowth (Figure 2E), and increased stem cell division at 24 hours post-feeding by only 13% (Figure 2F).

We cannot presently exclude the possibility that our defined diet formulations interfere with digestion or nutrient uptake. However, the small but significant impacts these food sources had on both animal size and neoblast division argue against complete malabsorption (Figure 2A-C). Furthermore, substitution of other thickening agents for xanthan gum in the complex diet had no discernable effect on the rate of degrowth (Table S1). Taken together, then, our results suggest total caloric intake is not the sole determinant of changes in animal size and stem cell division after feeding and are consistent with the possibility that liver contains unrecognized nutrients necessary for growth and capable of stimulating neoblast proliferation. In an effort to identify such hypothesized “missing ingredients,” we proceeded to screen over 40 different salts, vitamins, lipids, and other organic molecules, alone and in combination, for growth-promoting effects when added to the complex diet (Table S2). None had a visible impact. A definitive explanation for why *S. mediterranea* responds differently to consumption of oligidic and defined food sources, as well as development of a standardized laboratory diet capable of supporting long-term planarian maintenance, awaits further experimentation.

### Defined diets enable dsRNA delivery for gene knockdown

Planarians were one of the first animals in which gene expression was silenced by RNA interference after its discovery in *C. elegans*, and this technique has emerged as a powerful tool for studying gene function in numerous aspects of their biology (Fire et al., 1998; Sánchez Alvarado & Newmark, 1999; Reddien et al., 2005). dsRNA can be delivered via soaking or injection, or by feeding with either in vitro-transcribed dsRNA or dsRNA-expressing bacteria (Newmark et al., 2003; Orii et al., 2003; Rouhana et al., 2013; Adler & Sánchez Alvarado, 2018; Shibata & Agata, 2018). Many planarian investigators utilize RNAi feeding approaches because they combine the ease of soaking with the robustness of injection (Rouhana et al., 2013).

To assess the utility of the defined diets in RNAi feeding experiments, we compared their ability to induce previously documented phenotypes with that of the standard feeding approach (Materials and Methods). RNAi knockdown of the porphyrin biosynthesis enzyme *porphobilinogen deaminase-1 (PBGD-1)* prevents pigment biosynthesis in newly regenerated tissue (Stubenhaus et al., 2016). We found that a single RNAi feeding using either the complex or minimal diet was as effective in generating this phenotype as feeding with liver (Figure 3A). In order to compare phenotypic penetrance and expressivity, we next targeted the Wnt signaling component *β-catenin-1* and the exon junction complex subunit *magoh*. Knockdown of these genes results in the regeneration of two-headed animals from trunk fragments and stem cell loss in intact animals, respectively (Gurley et al., 2008; Petersen et al., 2008; Kimball et al., 2020). We did not observe a significant difference in the percentage of two-headed animals or the extent of stem cell depletion when comparing RNAi feeding for these genes using the complex diet versus feeding with liver (Figure 3B-D). These results demonstrate that both defined diets provide effective delivery mechanisms for dsRNA and reinforce the conclusion that ingested material is absorbed.

**Figure 3.**
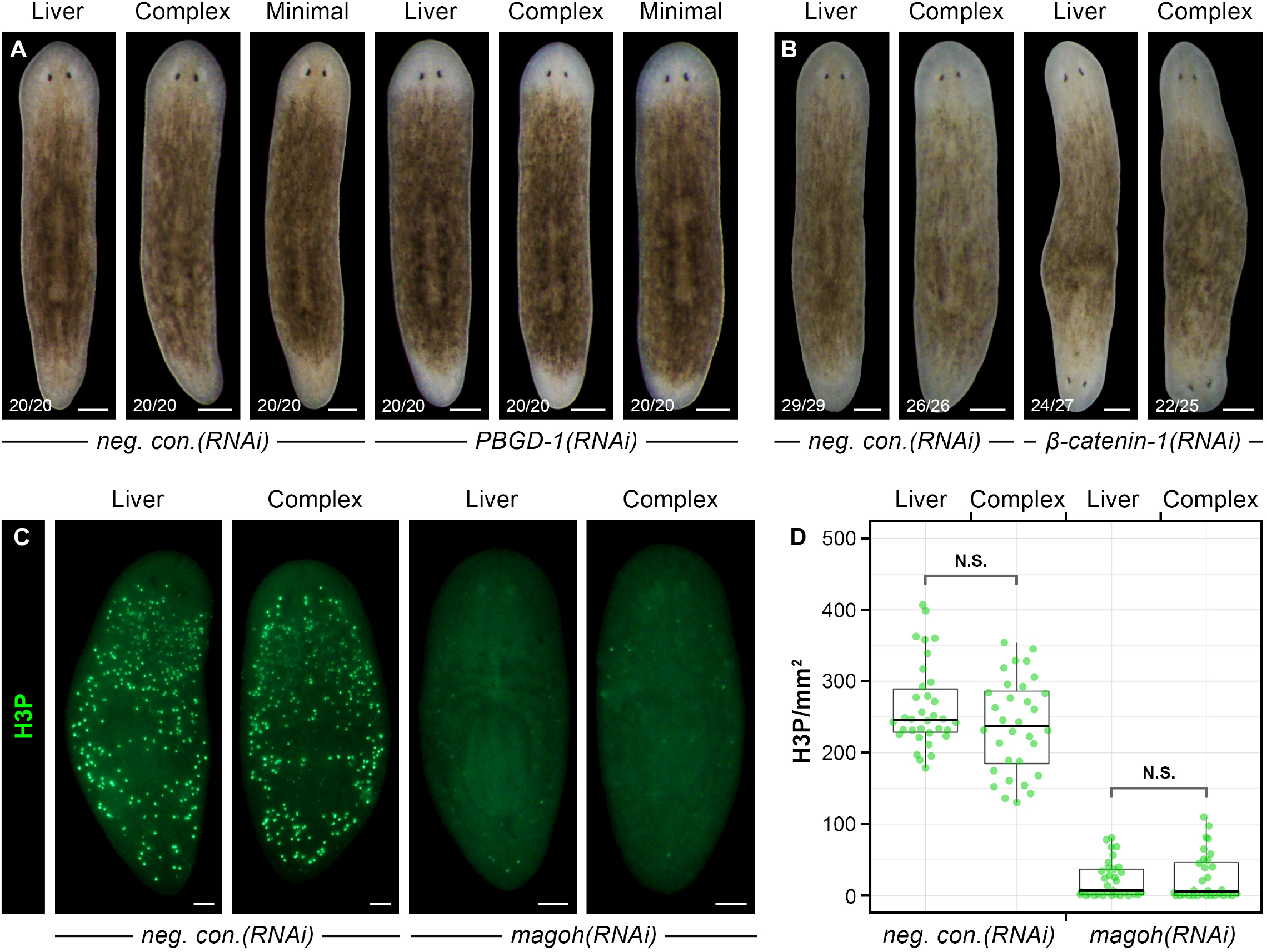
RNA interference by dsRNA delivery in defined diets. **(A,B)** Regardless of food source, RNAi knockdown of *PBGD-1* via a single dsRNA feeding (A) or of *β-catenin-1* via six dsRNA feedings (B) prevented pigmentation of newly formed tissue and resulted in two-headed animals, respectively. Images show representative live animals subjected to head and tail amputations the day after the final feeding and photographed at 21 days post-amputation. **(C,D)** H3P immunostaining in *negative control(RNAi)* and *magoh(RNAi)* animals fixed, labeled, and photographed three days after the last of four dsRNA feedings. The quantitative analysis was based on results from three biological replicates, with a minimum of 32 combined animals per condition. Boxes, whiskers, and horizontal lines denote the IQR, values within 1.5X the IQR, and medians, respectively. p values for T-tests comparing dsRNA delivery via liver and the complex diet were > 0.1 within each RNAi condition. Scale bars: (A,B) = 200 µm; (C) = 100 µm.

The ability to silence gene expression using the defined diets could offer important advantages over existing approaches in some experimental contexts. First, all previously described methods for dsRNA delivery provoke a more than two-fold increase in stem cell division as part of the normal responses to feeding (see above) or to injury (i.e. poke wounds produced by injection or amputations necessary for efficient gene knockdown by soaking) (Wenemoser & Reddien, 2010). Because the defined diets more modestly induce stem cell proliferation (Figure 2B,C), they provide an opportunity to achieve gene knockdown without as strongly impacting the homeostatic rate of neoblast division. Second, defined diet feeding allows for the easy introduction of dsRNA in the absence of contaminating bovine RNA (or DNA) from liver. This may be particularly advantageous in RNA-Seq experiments involving time points shortly after feeding. Third, the complex diet provides a means for rapid and repeated dsRNA delivery in the absence of organismal growth. This could be useful in experiments involving whole-mount labeling of RNAi animals by immunostaining, in situ hybridization, or TUNEL, all of which are typically more effective in smaller specimens (Brown & Pearson, 2015; Forsthoefel et al., 2018; Stubenhaus & Pellettieri, 2018). The above advantages may also apply to situations where delivery of other substances, such as nucleotide analogs used for stem cell labeling, is desired (Cheng & Sánchez Alvarado, 2018).

### Na azide treatment enables hydrogen peroxide bleaching of liver-fed animals

In the course of completing this line of research, we frequently encountered difficulties when attempting to fix liver-fed animals shortly after feeding. Specifically, we found that hydrogen peroxide used as a bleaching agent (Newmark & Sánchez Alvarado, 2000; Brown & Pearson, 2015; Forsthoefel et al., 2018) reacted with ingested liver to generate gas that damaged or destroyed specimens. We hypothesized this might be due to the high catalase activity present in liver and therefore tested whether the catalase inhibitor sodium azide might preserve tissue integrity during bleaching. Treatment with 1% sodium azide had the desired effect (Materials and Methods), eliminating visible gas production and allowing for H3P and in situ hybridization labeling in undamaged animals at one hour post-feeding (Figure 4). Animals fed the defined diets were effectively bleached and labeled without this modification.

**Figure 4.**
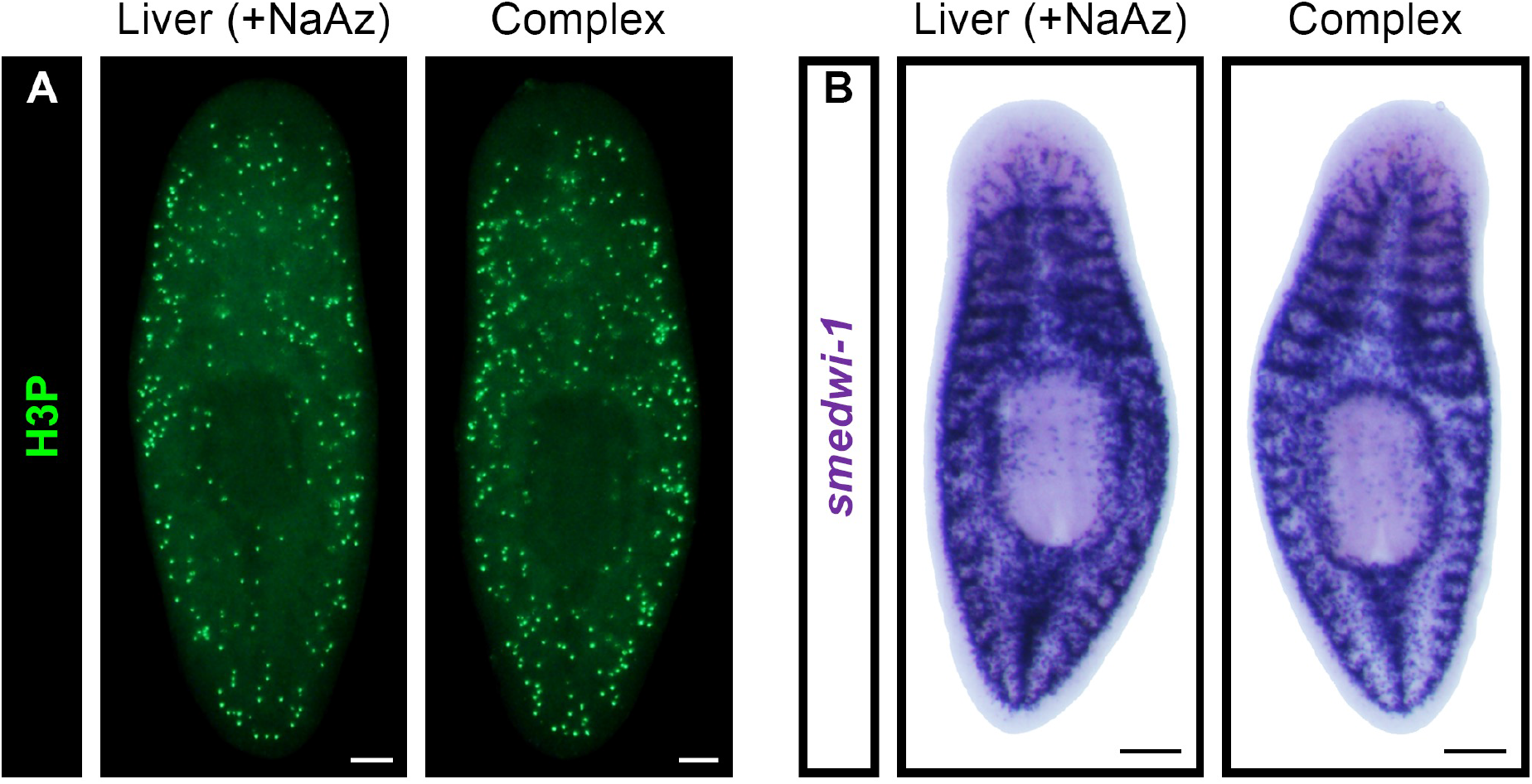
Labeling of recently fed animals. **(A,B)** H3P immunostaining (A) and whole-mount in situ hybridization with a probe for the neoblast marker *smedwi-1* (B) in animals fixed at one hour post-feeding. Liver-fed animals were pre-treated and bleached with Na azide (Materials and Methods). Scale bars: (A) = 100 µm; (B) = 200 µm.

## Conclusions

Nutritional studies in model organisms have historically focused on rodents, but experimentally accessible invertebrates such as *Drosophila* and *C. elegans* have recently assumed a more prominent role in the field. With their wealth of genetic tools, these systems are providing new mechanistic insight into topics such as feeding behavior, diet-disease interactions, and the links between metabolism and aging (Kapahi et al., 2017; Evangelakou et al., 2019; Zhou et al., 2019; Tierney, 2020). The development of partly or fully defined diets has played an important role in this work. As we have observed here, these diets are associated with lower rates of growth, development, and/or fecundity in comparison with oligidic food sources, limiting their utility in the long-term maintenance of laboratory populations (Lüersen et al., 2019; Zečić et al., 2019). Despite this drawback, they have proven extremely valuable in allowing for controlled manipulation of nutritional variables. We likewise expect the future application of these tools in planarians to bring about new advances in our understanding of the evolution and function of metabolic control mechanisms governing stem cell biology.

## MATERIALS AND METHODS

### Planarian maintenance

Clonal, asexual populations of *S. mediterranea* and *D. japonica*, as well as wild-caught *G. dorotocephala* (Carolina Biological), were maintained under standard laboratory conditions (Oviedo et al., 2008b; Accorsi et al., 2017; Merryman et al., 2018). Animals were propagated on a diet of homogenized calf liver with weekly or bi-weekly feedings and then starved for seven days prior to use in experiments (one month-starved animals were used for the long-term growth analysis in Figure 2A).

### Defined diet preparation and feeding

Complex and minimal diets were prepared according to the detailed protocols in Supplementary Materials and Methods, aliquoted, and stored at -20°C prior to use. Organic calf liver, used as a control, was homogenized as previously described (Merryman et al., 2018) and stored at -80°C. All experiments involved ad libitum feeding at room temperature – animals were allowed a minimum of one hour of unrestricted feeding in petri dishes (beyond this time point, feeding activity was no longer evident and unconsumed food invariably remained at the bottom of each dish). Fed animals were rinsed and returned to a 20°C incubator after feeding or processed and analyzed as indicated below. In all experiments involving multiple feedings, animals were fed every other day. Except for the long-term area measurements (Figure 2A) and analyses of relative food consumption (Figures 1C, 1D, and 2D), red or green food coloring was added to each diet (McCormick, Catalog No. 5210007107; 1 μl per 30 μl of food). This facilitated identification and removal of any animals that failed to eat (animals failing to eat were *not* removed for the experiments in Figures 1B, 1D, 2A, and 2D).

### Relative food consumption

Erioglaucine disodium salt (EDS) was mixed with each diet at a final concentration of 0.2% to enable a quantitative assessment of the relative amounts of food consumed (Ziman et al., 2020). Briefly, homogenates were prepared from five fed animals per condition in 100 μl of water by maceration with a pipet tip. Remaining tissue was pelleted in a microcentrifuge and absorbance measurements (620 nm) were obtained from supernatants in a BioTek Synergy HT plate reader. Negligible absorbance values for xanthan gum-only controls, which were not consumed, were subtracted from the readings for liver and the defined diets to account for any trace amounts of EDS absorbed via simple diffusion.

### Microscopy

All planarian photographs were obtained with an Olympus SZX16 stereomicroscope equipped with a DP72 digital camera. Cropped images of individual animals were placed on uniform black or white backgrounds to generate figure panels, with identical brightness and contrast adjustments applied within each experiment. Area measurements were made with Fiji/Image J (Schindelin et al., 2012).

### H3P immunostaining and WISH

H3P labeling/quantification and *smedwi-1* whole-mount in situ hybridization (WISH) were performed as previously described (Kimball et al., 2020). Boxplots displaying H3P results, as well as animal area measurements, were generated in R with PlotsOfData (Postma & Goedhart, 2019).

### RNA interference

RNAi knockdown experiments involved a standard feeding approach in which dsRNA-expressing bacteria were fed to animals in a mixture with the defined diets or homogenized calf liver (Newmark et al., 2003; Accorsi et al., 2017; Kimball et al., 2020). An equivalent amount of bacterial culture was used to prepare each RNAi food source. The *C. elegans* gene *unc-22* was employed as a negative control.

### Na azide treatment

Animals fixed within 24 hours of liver consumption were incubated overnight in 1% Na azide in PBST (1x PBS with 0.3% Triton-X), prior to bleaching overnight in 1% Na azide, 6% H_2_O_2_ in PBST. For WISH, these steps were followed by a formamide bleach without Na azide (Pearson et al., 2009; King et al., 2013).

## ACKNOWLEDGEMENTS

We thank J. P. Dustin, M. Ryan Woodcock, and Maggie Rice for assistance with preliminary experiments, and Shane Miller, Shane Merryman, and Alejandro Sánchez Alvarado for providing *S. mediterranea*.

## FUNDING

This work was supported by the National Science Foundation (Award No. 1656793) and by New Hampshire-INBRE through an Institutional Development Award (P20GM103506) from the National Institute of General Medical Sciences of the National Institutes of Health.

## COMPETING INTERESTS

The authors declare no competing or financial interests.

## AUTHOR CONTRIBUTIONS

Project conception and design: C.A., J.P.; acquisition of data: C.A., K.P., G.G.; analysis and interpretation of data: C.A., K.P., G.G., J.P.; writing the manuscript: J.P.

## TABLE AND FIGURE LEGENDS

**Figure S1.**
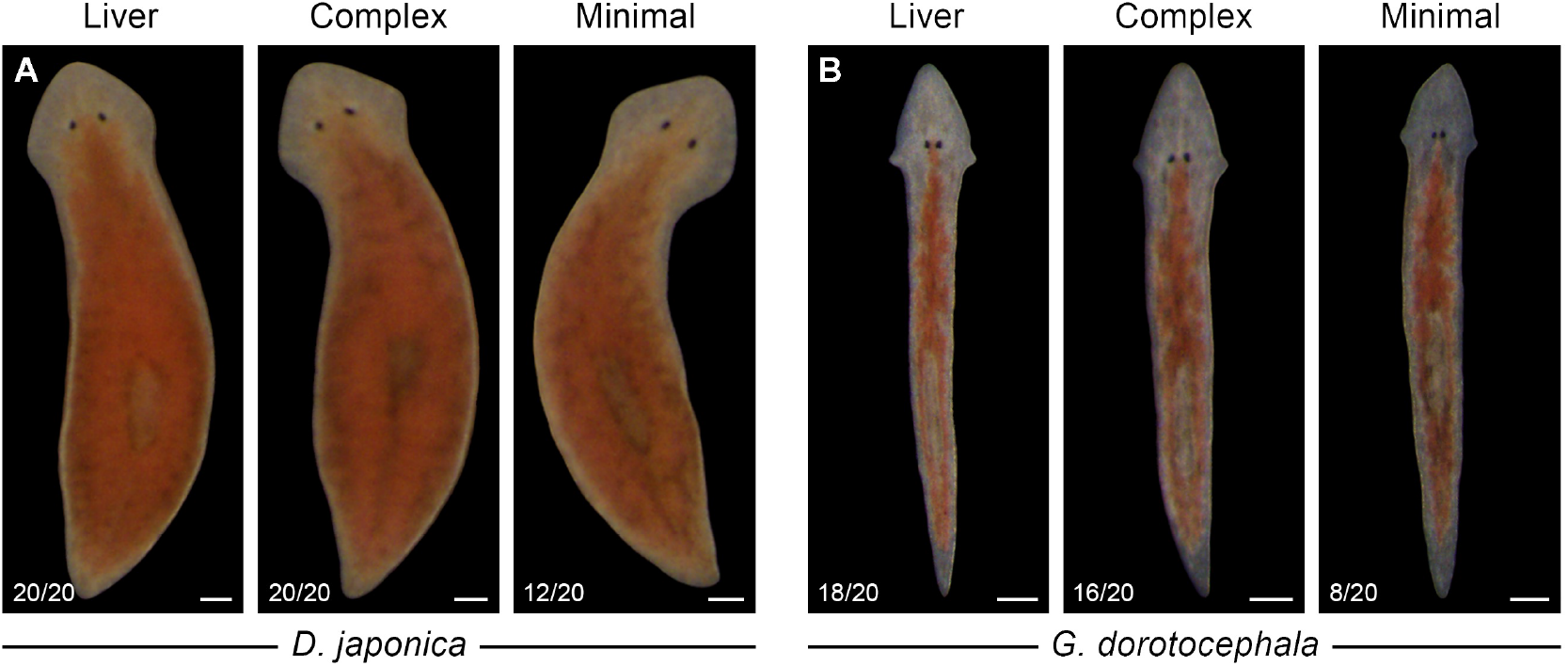
Defined diet feeding in *D. japonica* and *G. dorotocephala*. Live animals were photographed after a single ad libitum feeding with homogenized liver or defined diets. Red food coloring was added to visualize ingested food. The number of animals that consumed each diet is indicated in the lower left of each panel. Scale bars: 200 µm.

**Table S1.**
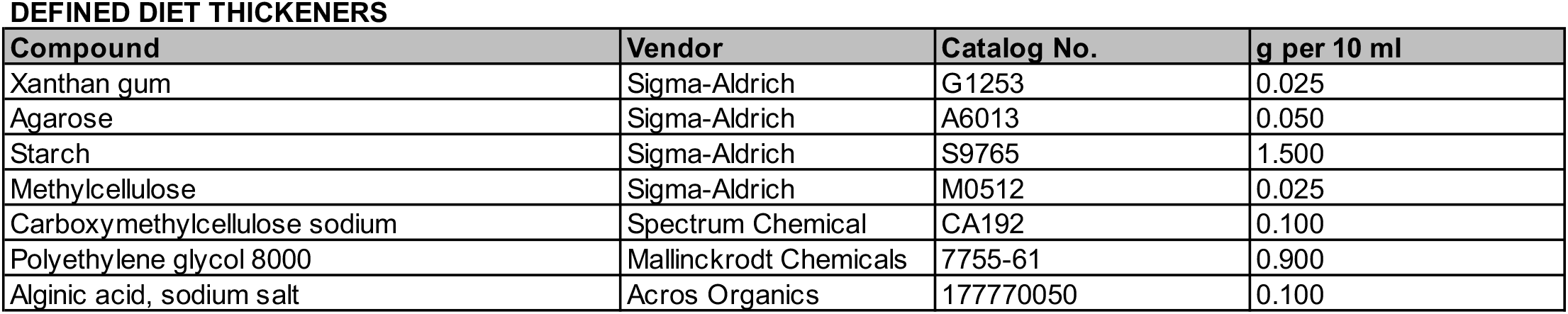
Alternative defined diet thickeners. Substitution of the indicated thickening agents for xanthan gum in the complex diet had no visible effect on the rate of degrowth (experiments were terminated when liver-fed controls demonstrated obvious growth).

**Table S2.** Tested complex diet supplements. Addition of the indicated compounds to the complex diet, alone or in combination, had no visible effect on the rate of degrowth (experiments were terminated when liver-fed controls demonstrated obvious growth).

| **Compound** | **Vendor** | **Catalog No.** | **g per 100 ml** |
| --- | --- | --- | --- |
| Potassium phosphate monobasic | Fisher Scientific | BP362 | 0.0500 |
| Magnesium sulfate heptahydrate | Sigma-Aldrich | 230391 | 0.0100 |
| Sodium chloride | Bio Basic | SB0476 | 0.0010 |
| Calcium chloride dihydrate | Bio Basic | CD0050 | 0.0010 |
| Ammonium iron (III) sulfate dodecahydrate | Acros Organics | 205881000 | 0.0010 |
| Copper (II) chloride dihydrate | Acros Organics | 405840050 | 0.0010 |
| Manganese (II) chloride tetrahydrate | Sigma-Aldrich | M3634 | 0.0010 |
| Zinc chloride | Acros Organics | 196840050 | 0.0010 |
| L-Ascorbic acid | Fisher Scientific | A61 | 0.0100 |
| D-Calcium pantothenate | Acros Organics | 243300050 | 0.0010 |
| Cyanocobalamin | Alfa Aesar | A14894 | 0.0010 |
| Adenosylcobalamin | Cayman Chemical | 21571 | 0.0010 |
| Pyridoxine hydrochloride | Alfa Aesar | A12041 | 0.0010 |
| Riboflavin 5' phosphate sodium dihydrate | Alfa Aesar | J66949 | 0.0010 |
| Thiamine hydrochloride | Acros Organics | 148990100 | 0.0010 |
| Nicotinic acid | Acros Organics | 128291000 | 0.0010 |
| Lipoic acid | Acros Organics | 138720050 | 0.0010 |
| D-Biotin | Acros Organics | 230090010 | 0.0010 |
| Folic acid | Acros Organics | 216630100 | 0.0010 |
| Beta carotene | Sigma-Aldrich | C9750 | 0.0010 |
| Retinol acetate | MP Biomedicals | 103257 | 0.0010 |
| Cholecalciferol | Alfa Aesar | B22524 | 0.0001 |
| Palmitic acid | TCI America | P0002 | 0.0010 |
| Stearic acid | TCI America | S0163 | 0.0010 |
| Cholic acid | Chem-Impex | 00071 | 0.0010 |
| Cholesterol | Sigma-Aldrich | C8667 | 0.0010 |
| RNA from torula yeast | Sigma-Aldrich | R6625 | 0.0200 |
| DNA, sodium salt from salmon sperm | GoldBio | D-112 | 0.0100 |
| Betaine | Alfa Aesar | B24397 | 0.0100 |
| L-Carnitine | Acros Organics | 241040010 | 0.0100 |
| N-Acetylglucosamine | Alfa Aesar | A13047 | 0.0100 |
| Choline chloride | Acros Organics | 110295000 | 0.0050 |
| Myo-inositol | Sigma-Aldrich | I7508 | 0.0050 |
| Cytochrome C | Sigma-Aldrich | C2506 | 0.0010 |
| Hemin chloride | Sigma-Aldrich | 3741 | 0.0001 |
| DL-Alpha tocopherol | Santa Cruz Biotechnology | sc-294383 | 0.0095 |
| Alpha linoleic acid | Cayman Chemical | 90210 | 0.0075 |
| Oleic acid | Alfa Aesar | 31997 | 0.0089 |
| Linoleic acid | Beantown Chemical | 129310 | 0.0090 |
| Selenium | Ultra Scientific | IAA-234 | 0.0005 |

## SUPPLEMENTARY MATERIALS AND METHODS

### Complex Diet Preparation

1. **Prepare AA#1 Solution**
  a. Add to a 100 ml beaker:
    - 0.32 g L-Valine
    - 0.34 g L-Alanine
    - 0.48 g L-Leucine
    - 0.14 g L-Methionine
    - 0.24 g L-Isoleucine
    - 0.09 g L-Cysteine
  b. Add RO water to 50 ml and stir until amino acids dissolve. Store at 4°C.
2. **Prepare AA#2 Solution**
  a. Add 0.05 g L-Tyrosine to a 100 ml beaker.
  b. Add ∼40 ml RO water and bring to a boil.
  c. Stir to dissolve and bring to final volume of 50 ml with RO water.
  d. Bring to room temperature before using for food preparation.
  e. Use fresh solution each time food is prepared.
3. **Prepare AA#3 Solution**
  a. Add to a 100 ml beaker:
    - 0.22 g L-Phenylalanine
    - 0.17 g L-Threonine
    - 0.41 g L-Aspartic acid
    - 0.70 g L-Glutamic acid monosodium salt hydrate
    - 0.35 g Glycine
    - 0.25 g L-Proline
    - 0.20 g L-Serine
    - 0.06 g L-Glutamine
    - 0.13 g L-Histidine
    - 0.41 g L-Lysine monohydrochloride
    - 0.34 g L-Arginine monohydrochloride
    - 0.05 g L-Tryptophan
    - 0.04 g L-Asparagine
  b. Add RO water to 50 ml and stir until amino acids dissolve. Store at 4°C.
4. **Prepare Dextrose Solution**
  a. Add 21.70 g dextrose to a 100 ml beaker.
  b. Add ∼40 ml RO water and bring to a boil.
  c. Stir to dissolve and bring to final volume of 50 ml with RO water.
  d. Bring to room temperature before using for food preparation. Store at 4°C.
5. **Prepare Xanthan Gum Solution\***
  a. Add 0.25 g xanthan gum to a 15 ml conical tube.
  b. Add RO water to 15 ml and vortex thoroughly to mix. * Note: other thickeners can be substituted if desired (see Table S2).
6. **Prepare Food**
  a. Add to a 50 ml conical tube:
    - 0.70 g lecithin (use waxed weighing paper)
    - 2,316 μl AA#1 Solution
    - 1,000 μl AA#2 Solution
    - 3,184 μl AA#3 Solution
    - 50 μl Dextrose Solution
  b. Vortex thoroughly to mix.
  c. Add 0.30 g glycogen and 0.5 ml RO water.
  d. Vortex thoroughly to mix.
  e. Adjust pH to 6.9-7.0 with 1M NaOH.
  f. Add 1.5 ml xanthan gum solution.
  g. Bring to final volume of 10 ml with RO water.
  h. Vortex thoroughly to mix. Aliquot and store at -20°C.

### Minimal Diet Preparation

1. **Prepare Xanthan Gum Solution\***
  a. Add 0.25 g xanthan gum to a 15 ml conical tube.
  b. Add 15 ml RO water and vortex thoroughly to mix. * Note: other thickeners can be substituted if desired (see Table S2).
2. **Prepare Food**
  a. Add 0.70 g lecithin to a 50 ml conical tube (use waxed weighing paper).
  b. Add 8 ml RO water and vortex thoroughly to mix.
  c. Adjust pH to 6.9-7.0 with 1M NaOH.
  d. Add 1.5 ml xanthan gum solution.
  e. Bring to final volume of 10 ml with RO water.
  f. Vortex thoroughly to mix. Aliquot and store at -20°C.

## Notes

### Competing Interest Statement

The authors have declared no competing interest.

